# EZH2 Inhibition Boosts Tumor Immunogenicity and TIL Reactivity in Lung Cancer

**DOI:** 10.1101/2025.07.12.663845

**Authors:** Daniel R. Plaugher, Christian M. Gosser, Yindan Lin, Avery R. Childress, Jinpeng Liu, Jinze Liu, Xufeng Qu, Tanner DuCote, Dave-Preston Esoe, Siva K Gandhapudi, Christine F. Brainson

**Affiliations:** Department of Toxicology and Cancer Biology, University of Kentucky, Lexington KY; Department of Cancer Biostatistics, University of Kentucky, Lexington KY; Massey Comprehensive Cancer Center, Virginia Commonwealth University, Richmond, VA; Department of Biostatistics, Virginia Commonwealth University, Richmond, VA; Department of Microbiology, Immunology & Molecular Gen, University of Kentucky, Lexington KY; Markey Cancer Center, University of Kentucky, Lexington KY

**Keywords:** non-small cell lung cancer, cell signaling, computational biology, immune micro-environment, epigenetics, tumor-infiltrating lymphocytes

## Abstract

Lung cancer, the leading cause of cancer-related death in the United States, remains a significant public health burden. One of the most personalized treatments to date uses a patient’s own tumor-infiltrating lymphocytes (TILs) as a cellular therapy to target the tumor. Endogenous TIL activity is also essential for immune checkpoint inhibitor (ICI) effectiveness. However, responses to immunotherapies vary due to immune-suppressive tumor microenvironments (TMEs) and limited antigen presentation. In this study, we computationally examine cell-cell signaling and transcriptional activity using single-cell RNA sequencing of lung cancer treated by inhibiting methyltransferase EZH2. We show that EZH2 inhibition shifts the TME to immunogenic signaling patterns conducive to increased T cell response, including antigen presentation and homing. T cells also showed more stem-like phenotypes. Importantly, inferred EZH2 activity showed it can still perform non-methyltransferase duties that are vital for T cell differentiation. Lastly, we tested our findings with patient-derived NSCLC TILs and found that transient EZH1/2 inhibition with valemetostat prior to harvest increases the CD8:CD4 ratio and T cell reactivity, but preferentially expands effector memory rather than central memory populations. These results indicate that transitory EZH2 inhibition could improve immunotherapies for lung cancer patients, with additional optimization needed to maximize TIL therapeutic benefit.

## Main

Lung cancer is the leading cause of cancer-related death in the U.S. among both men and women. With a combined 131,889 deaths in 2022, lung cancer’s closest competitor for total deaths is colorectal cancers at 52,967 deaths(^1^). While survival rates have slowly climbed due to advances in therapy and early detection, the 5-year survival rate remains at 27% for all stages with lower rates for advanced-stage disease. Lung cancer is well known for its immense heterogeneity, leading to limited effective treatment options. However, there is a silver lining; NSCLC is highly mutated, and this is a vulnerability due to the high rate of neo-antigen presentation for T cell recognition(^2, 3^).

Novel immunotherapeutic techniques including anti-PD/PD-L1 immune checkpoint inhibitor (ICIs) and adoptive cell transfer (ACT) have emerged as effective therapies for many cancer types. While anti-PD1/PD-L1 antibodies rely on endogenous T cell activities, ACT is a “living” treatment where lymphocytes are infused into a patient and continue to expand after administration (more than 1000-fold)(^2^). Chimeric antigen receptor T cell (CAR-T) and modified T cell receptor (TCR) therapies are designed to recognize specific (neo)antigens, an approach that has fallen short in solid tumors due to lack of common antigens among patients(^4^). Tumor-infiltrating lymphocyte (TIL) therapy is polyclonal by nature and is arguably the most personalized form of cancer therapy. With respect to NSCLC, it is currently unknown what tumor genotypes and phenotypes will respond best to TIL therapy, but numerous trials combining ICIs and TIL therapy are underway – including studies investigating epigenetic modulation(^4^).

A key challenge for both TIL and ICI therapies is to maintain T cell stemness and resist exhaustion, while also ensuring the patient’s tumor microenvironment (TME) is immunogenic. These phenotypes can be controlled through epigenetic mechanisms. For example, DNA and histone methylation are typically associated with gene silencing(^5^). Indeed, multiple studies over decades of research have shown the importance of EZH2 (enhancer of zeste homolog 2) in its role as the catalytic subunit of the Polycomb Repressive Complex 2 (PRC2), where it tri-methylates histone 3 at lysine 27 (H3K27me3) through its SET domain. Therapeutic options targeting EZH2 inhibition (EZH2i) include, but are not limited to, GSK126(^6^), tazemetostat(^7, 8^), and valemetostat(^9^) (a dual EZH1/2 inhibitor. An incredibly important point to note is that these inhibitors do not act as knock-out drugs but rather are specifically geared to target the SET domain and could still allow non-methyltransferase activities described below. This distinction is key because studies have shown that *deletion* of *Ezh2* can restrict memory T cell function and drive terminal differentiation (^10, 11^). However, other studies have shown that *Ezh2 deletion* can reprogram regulatory T cells (Treg) to T-effector cells (T_EFF_), and lead to de-repression of transcription factors *Tcf7* and *Id3* important for memory T cells (T_M_) (^10, 12–14^).

Even though EZH2 is not known to be a DNA binding transcription factor (TF), beyond its canonical role catalyzing H3K27me3 within PRC2, EZH2 is reported to exert non-canonical functions that reshape transcriptional and signaling networks(^15^). In a PRC2-dependent manner, EZH2 can methylate non-histone substrates such as GATA4, repressing acetylation and gene activation (^16^). However, EZH2 acts independently of PRC2 as a co-activator or enzyme that modifies targets. In prostate cancer, EZH2 directly occupies the androgen receptor (AR) locus to activate transcription without requiring SUZ12, EED, or H3K27me3 (^17^). Similarly, EZH2 is independently associated with ERα–β-catenin, NF-κB (RelA/RelB), Trim28–SWI/SNF, MYC, E2F1, or Pol II to drive oncogenic programs such as c-MYC, CCND1, IL6/TNF, and IGF1R expression(^18^). It has been shown that phosphorylation by AKT1, JAK3, and p38α disengage EZH2 from PRC2, permitting it to function independently of PRC2 to methylate (when in a complex with AR) non-histone substrates like STAT3 or RORα, and/or to serve non-enzymatic roles in mRNA translation and DNA-damage repair(^16, 18^). Collectively, these PRC2-dependent non-histone and PRC2-independent mechanisms illustrate EZH2’s complex identity as both epigenetic silencer and transcriptional activator.

Recently, multiple groups have shown that EZH2i enhances CAR-T, TCR, and ICI therapies. Specifically, Hou et. al.(^19^) demonstrated that transient EZH2i promotes progenitor-like T cells without compromising cell expansion. They further revealed that EZH2i improves *in vivo* proliferation, homeostasis, and recall response of ACT T cells, while also showing that there were less-differentiated T cells in human PBMCs and that CAR T-cell exhaustion was alleviated. Then subsequently Porazzi et al. (^20^) showed similar results such that EZH2 and EZH1/EZH2 inhibition improves the anti-tumor efficacy of CAR-T- and TCR T cell-based therapies against multiple liquid and solid tumors. In our own study(^21^), we demonstrated that EZH2i can increase ICI (anti-PD1) response in lung squamous cell carcinomas (LSCCs) using tazemetostat and GSK126 as a tool compound. Our *in vitro* experiments using 2D human cancer cell lines, and 3D murine and patient-derived organoids all showed that EZH2i leads to up-regulation of both major histocompatibility complex class I and II (MHCI/II) expression at mRNA and protein levels. We then used ChIP-sequencing to confirm the loss of EZH2-mediated histone marks, as well as the gain of activating histone marks at key loci. Within murine autochthonous and syngeneic LSCC models, we demonstrated robust tumor control with ICI+EZH2i combination therapy – with significant decreases in tumor cells, as well as tumor promoting macrophage and neutrophil populations. In contrast, we saw significant increases in T cells, cycling cells, normal lung, and some anti-tumor neutrophil populations(^21^).

Despite these advances, our prior study largely focused on myeloid compartments and lacked a systems-level view of how the immune TME changes under EZH2 inhibition. Here, we performed new downstream analysis of single-cell RNA-sequencing (scRNAseq) data from lung tumors and total lung of our autochthonous murine model (Figure-1A). Using a top-down framework, we began with global TME cell-cell signaling, then narrowed to pathway-level signaling, and ultimately resolved to specific cell-type activity to understand how EZH2 inhibition remodels the TME to support T cell responses (Figure-1B). We first present inferred changes in cell-cell communication using CellChat(^22^) and LIANA(^23^), with a primary focus on endogenous TILs (eTILs) that have not undergone *ex vivo* expansion. We then investigated changes in TF activity using tools like SCENIC(^24^), BITFAM(^25^), and CollecTRI(^26^). Finally, we validated our key observations by applying EZH1/2i (valemetostat) to human NSCLC TILs that have undergone the full TIL production process. These insights are critical for determining whether EZH2 inhibition can effectively augment T cell-based immunotherapies for patients with lung cancer and other malignancies.

**Fig. 1:**
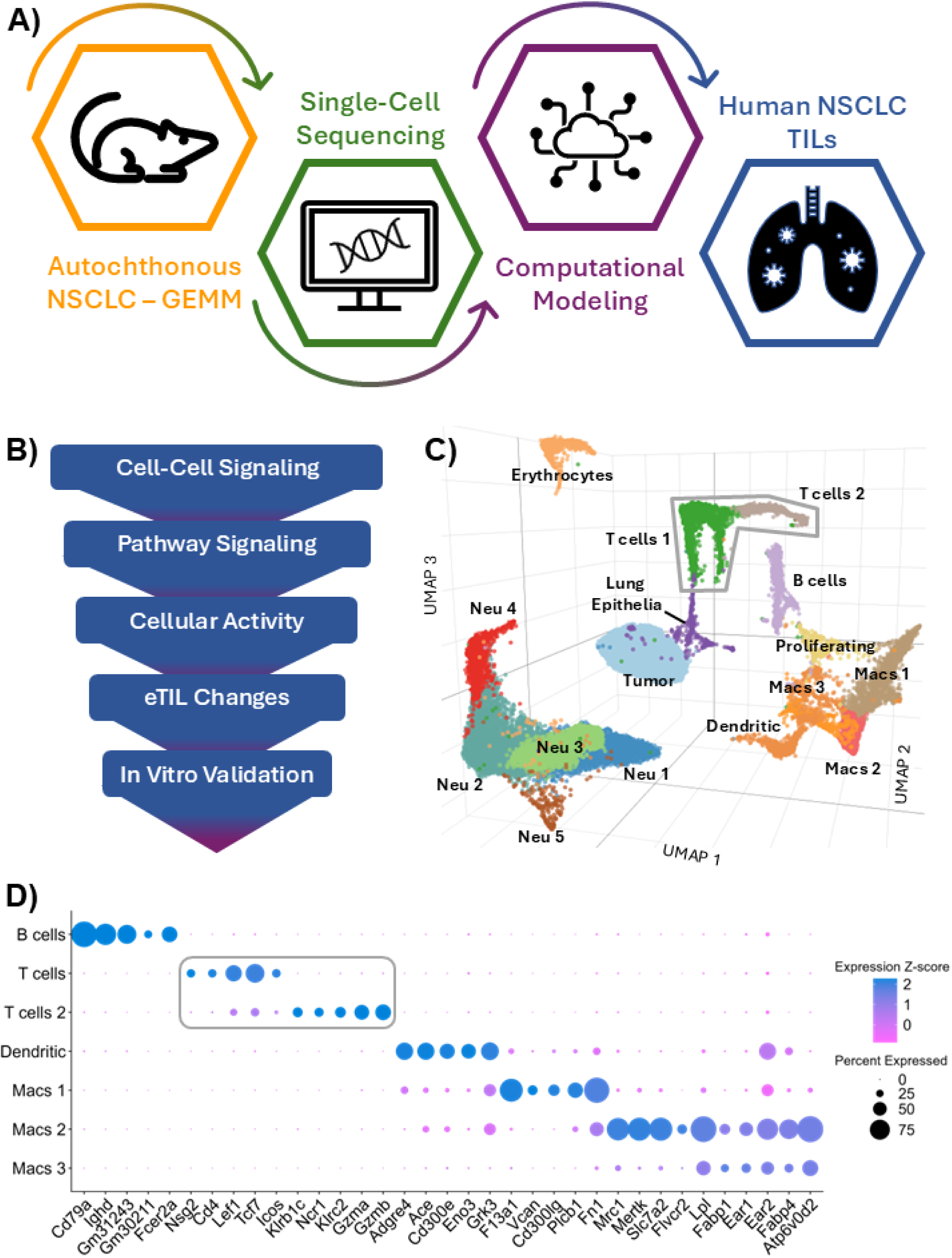
Single-cell profiling of lung TME. **(A)** Schematic of workflow. **(B)** Framework of top-down approach. **(C)** Annotated 3D UMAP showing 16 populations within lung tumors with concatenated populations of Veh-PM, EZH2i-PM, Veh-M, and EZH2i-M. **(D)** Dot plot for top 5 defining markers among lymphocytes and classic antigen-presenting cells, according to expression strength and normalized z-score. Abbreviations: Neu – neutrophil; Macs – macrophage; Prol. – proliferating; Lung Ep – lung epithelia

## Single cell landscape

Fresh autochthonous lung tumors (malignant - M) were dissociated and analyzed via single-cell RNA sequencing after four weeks of treatment with GSK126 (Veh-M , EZH2i-M), we used premalignant (PM) total lung treated for two weeks as additional controls (Veh-PM, EZH2i-PM). Throughout our downstream analysis, we focus on pairwise changes among these conditions. Overall, we identified 16 unique cell populations that were annotated based on conserved markers and plotted in a 3D UMAP (Figure-1C**; Supp. Figure-1A**). Markers for lymphocytes and traditional antigen-presenting cells are highlighted in Figure-1D, and cell-type proportions with differential markers can be found in **Supplementary** material. Specifically, among clusters of endogenous TILs, Tcells1 exhibit more progenitor and memory-like markers (*Il7r, Ccr7, S1pr1, Tcf7, Lef1*), whereas Tcells2 represent more effector and exhausted phenotypes (*Gzmb, Ifng, Ccl5, Nkg7, Klrg1*).

## Cell-cell signaling patterns shift after EZH2 inhibition

For a broad view of cellular communications, we employed CellChat(^22^) to infer interactions (and strength of interaction) among ligands-receptor pairs and their cofactors, with LIANA(^23^) as a supporting method (see **Supplementary**). We first observed that EZH2i treatment resulted in minimal changes in premalignant overall interactions (Veh: 3,086 vs. EZH2i: 3,059; 0.9% reduction) and interaction strength (Veh: 111.3 vs. EZH2i: 91.72; 18% reduction). In contrast, EZH2i in malignant samples led to substantial reductions in both the number of interactions (Veh: 3,057 vs. EZH2i: 2,231; 27% reduction) and interaction strength (Veh: 97.6 vs. EZH2i: 63.5; 35% reduction) (**Supp. Figure-1B**). A deeper dive using differential heatmaps reveals explicitly which cell-cell signals are changing, where red (or blue) represents increased (or decreased) signaling in treatment compared to vehicle (Figure-2A). The top-colored bar plot represents the column sum of values displayed in the heatmap (incoming signaling) and right colored bar plots represent the sum of row of values (outgoing signaling).

**Fig. 2:**
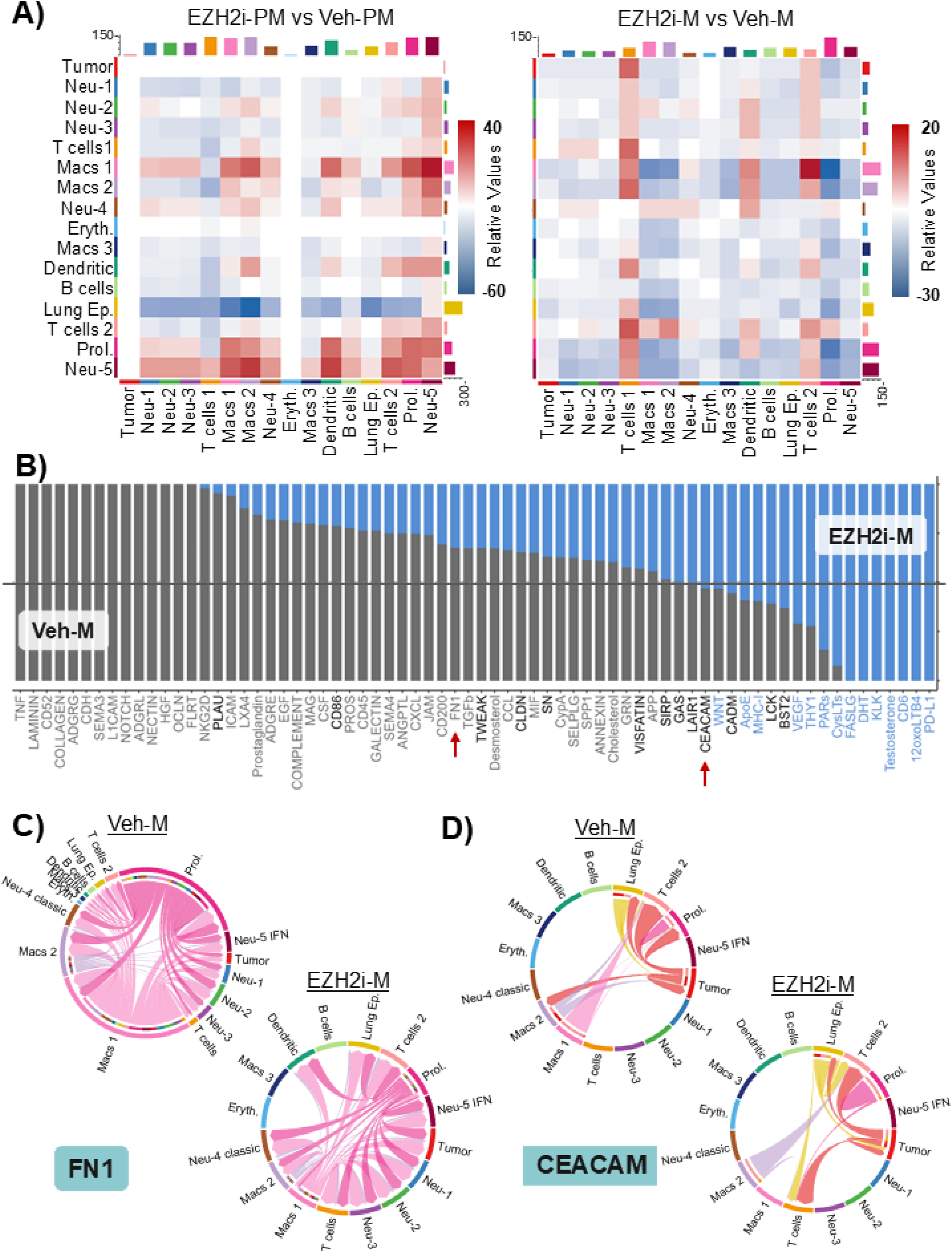
Cell-cell communication patterns. **(A)** Differential interaction heatmaps, where red (or blue) represents increased (or decreased) signaling in treatment compared to vehicle. The top-colored bar plot represents the column sum of values displayed in the heatmap (incoming signaling) and right colored bar plots represent the sum of row of values (outgoing signaling). **(B)** Conserved and context-specific relative information flow alterations for highly significant (p<0.01) pathways. Pathways with gray text are enriched in Veh-M, blue is enriched in EZH2i-M, and black indicates an enriched pathway in both groups. Red arrows are markers for pathways highlighted in remaining sub-figures. **(C&D)** Chord diagrams showing signaling for CEACAM and FN1 (respectively) between cell groups, where inner thinner bar colors represent the targets that receive signal from the corresponding outer bar. Inner bar size is proportional to the signal strength received by the targets. Abbreviations: Neu – neutrophil; Macs – macrophage; Prol. – proliferating; Lung Ep – lung epithelia

When comparing EZH2i-PM versus Veh-PM, we generally see a decrease in most neutrophil signaling, but increases in macrophage (1&2), dendritic, proliferating, and Neu-5. Importantly, we previously showed that Neu-1, Neu-4, and Neu-5 express genes such as *Tnf*, *Cxcl10*, and numerous IFN-response genes(^21^). Tcells1 only has increased signaling with Neu-5 and decreases elsewhere, but Tcells2 has increased communication with anti-tumor neutrophils, proliferating cells, autocrine signaling, Macs 1, and dendritic cells. In contrast, differential signaling of EZH2i-M versus Veh-M nearly all decreases with the interesting exception of eTILs and dendritic cells. We see very distinct increases in outgoing signals of Tcells1, Tcells2, and dendritic cells, and perhaps most importantly there is increased tumor signaling unique to only eTILs (CEACAM, MHC-I, CXCL, KLK, and ADGRE pathways; **Supp. Figure-1C**). Collectively, the enhanced communication between antigen-presenting cells, IFN-responsive neutrophils, and T cell populations indicates that EZH2 inhibition reprograms the TME to be more immunogenic through increased immune activation and antigen-presenting capacity.

Next, we dove deeper to identify conserved and context-specific signaling pathways among all cell types by comparing information flow, determined by the sum of communication probabilities among all pairs of cell groups (the total weights in the network). Highly significant signaling pathways (p<0.01) were ranked based on differences in the overall information flow between treatment groups and presented in a stacked bar graph (Figure-2B). The top signaling pathways colored gray (text) are enriched in Veh-M, blue text is enriched in EZH2i-M, and black text indicates an enriched pathway in both groups. We can identify signaling pathways that turn off after treatment (all gray), decrease (majority gray), turn on (all blue), or increase (majority blue) by changes to their information flow when comparing conditions. First, it is apparent that EZH2i results in fewer enriched pathways (14) compared to Veh-M (43). There is a loss of 14 total pathway signals (e.g. CD52, COLLAGEN, ADGRG, NOTCH, LAMININ…etc.) and 7 total signals gained (FASLG, KLK, CD6…etc.) when treatment is applied. We also see a loss (or decrease) of many pathogenic pathways after treatment, such as collagen and fibronectin that play major roles in pulmonary fibrosis(^27^). Using a chord diagram, we show that fibronectin signals via FN1 are decreased and much weaker after treatment (Figure-2C). Further, there are increases in immune infiltration pathways like CEACAM, MHC-I, LCK, and THY1. The CEACAM pathway has been shown to positively correlate with immune infiltration(^28^), and in Figure-2D we see that immune signaling increases primarily to eTILs from the tumor and lung epithelium. Taken together, all these results indicate a shift in the immune TME from suppressive to permissive.

## Ezh2 signatures are maintained despite methyltransferase inhibition

To determine if SET domain inhibitors completely shut down EZH2 activity, we used the SCENIC(^24^) pipeline to infer transcription modification activity between treatment groups, with BITFAM(^25^) and CollecTRI(^26^) as supporting approaches (see **Supplementary**). Note that some “TFs” in the prebuilt SCENIC database are more accurately identified as modifiers (e.g. *Dnmt1, H2faz, Ezh2 …etc.)* but will still be referred to as TFs for ease.

Further, gene expression level does not imply activity level(^24^),and while EZH2 is not a known DNA binding TF, it is known to have PRC2-independent activity as discussed in the introduction(^16–18^). Between all three inference techniques, we normalized and averaged total EZH2 activity in Figure-3A. We see little fluctuation within pairwise comparisons; however, it is evident that the malignant environment significantly impacts the signatures of EZH2.

**Fig. 3:**
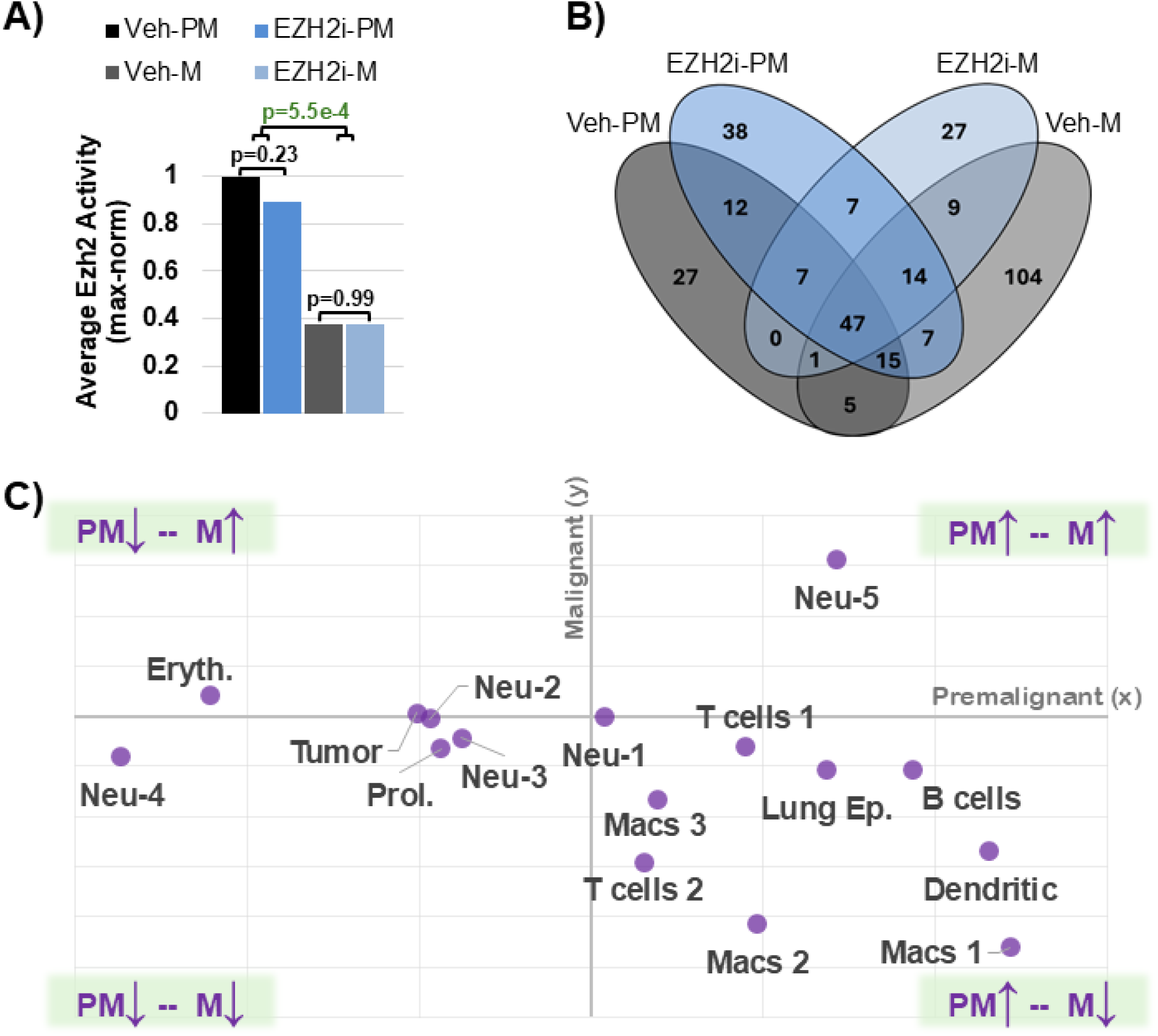
Computationally inferred transcription activity. **A)** Levels of inferred Ezh2 activity (max-normalized) using pySCENIC, BITFAM, and CollecTRI inference methods. **B)** 4-way Venn diagram of inferred Ezh2 target totals. A precise list for each treatment group is available on GitHub. **C)** Scatter plot of cell-type specific Ezh2 activity log2FC between PM (x-axis) and M (y-axis). Abbreviations: Neu – neutrophil; Macs – macrophage; Prol. – proliferating; Lung Ep – lung epithelia

Overall, SCENIC showed that EZH2 had 114 inferred targets in Veh-PM, 147 in EZH2-PM, 202 in Veh-M, and 112 in EZH2i-M (see *EZH2_total_targets.txt* on GitHub). We show how these targets overlap with a 4-way Venn diagram (Figure-3B). Interestingly, the treatment groups share 7 unique inferred targets that encompass cell cycle control (*Cks2, Hist1h3d*), metabolic transport (*Slc7a2, Far1*), ER function (*Calu*), and migration-related signaling (*Il11ra1, Nav2*) - indicating possible processes targeted by EZH2i.

Between cell types and across both PM and M, Neu-5 cells show the largest log_2_FC increase in EZH2 activity post-EZH2i, and Neu-4 cells have the greatest decrease (Figure-3C). For discordant outcomes between PM and M, erythrocytes exhibit the largest increase from PM to M, while Macs1 show the greatest decrease. Within eTIL populations, activity in Tcells1 increases modestly in PM (+0.90) but decreases in M (−0.62), and Tcells2 showed a smaller increase in PM (+0.31) followed by a stronger decrease in M (−2.93). This suggests that EZH2i may modulate already low EZH2 expressing TILs enough to shift some effector-like cells toward a more terminally exhausted state, consistent with the higher susceptibility of Tcells2 that are enriched for effector/exhausted phenotypes. However, to the contrary, CollecTRI analysis instead indicates that EZH2 activity in eTILs is elevated relative to vehicle (Tcells1: +42% PM, +40% M; Tcells2: +75% PM, +38% M; see **Supplementary**). Likewise, BITFAM shows that aggregate eTILs see a modest EZH2 activity increase (PM: +7.8%; M: +3.7%; see **Supplementary**). The disagreement across methods suggests that EZH2 activity in eTILs lies close to a threshold where modest shifts can be interpreted differently, highlighting the challenge of resolving EZH2i responses in these low-signal states. However, all methods crucially show that EZH2 activity is never fully lost.

Complementary analysis (see **Supplementary; Supp. Figures-2&3**) includes Regulon Specificity Score (RSS) that calculates cell type-specific regulon activity, regulon size and target comparisons, regulon correlation. We also infer gene regulatory networks (GRNs) and performed network analysis for connectivity, connected components (weak and strong), structure, centrality, motifs, and targets for phenotype control(^29^).

## Endogenous TIL signaling landscape is reprogrammed after EZH2 inhibition

As a continued point of distinction, here “eTILs” are referencing the endogenous infiltrates that we have subset from our primary population (Figure-1C). The first notable observation is the raw count variability between eTILs exposed to pre-malignant and malignant TMEs: pre-malignant samples (Veh-PM/EZH2-PM) show ∼20-fold higher counts than malignant samples (**Supp. Figure-1A**), consistent with the well-recognized immune-suppressive TME of LSCC. Among malignant samples, however, EZH2i-M exhibits the highest proportion of *Cd8*+ eTILs (Figure-4A) suggesting that EZH2i may promote proportionally more cytotoxic TIL infiltrates. In contrast, *Cd4*+ eTILs are predominantly seen in premalignancy but drop the lowest in EZH2i-M. Accordingly, the *Cd8*+:*Cd4*+ ratio is greatest in EZH2i-M, indicating a strong shift to more cytotoxic T cell populations. As previously noted, Tcells1 aligns more with stem-like phenotypes compared to Tcells2. To complement these observations, we performed trajectory analysis to assess differentiation and pseudotime, seeding the graph at the *Tcf7*-high region as the root node (**SUPP. FIGURE-4).** Notably, proportions in Figure-4A do not include *Cd3*+/*Cd8*-/*Cd4*- (double negative) populations such as γδ-T or NK-T.

**Fig. 4:**
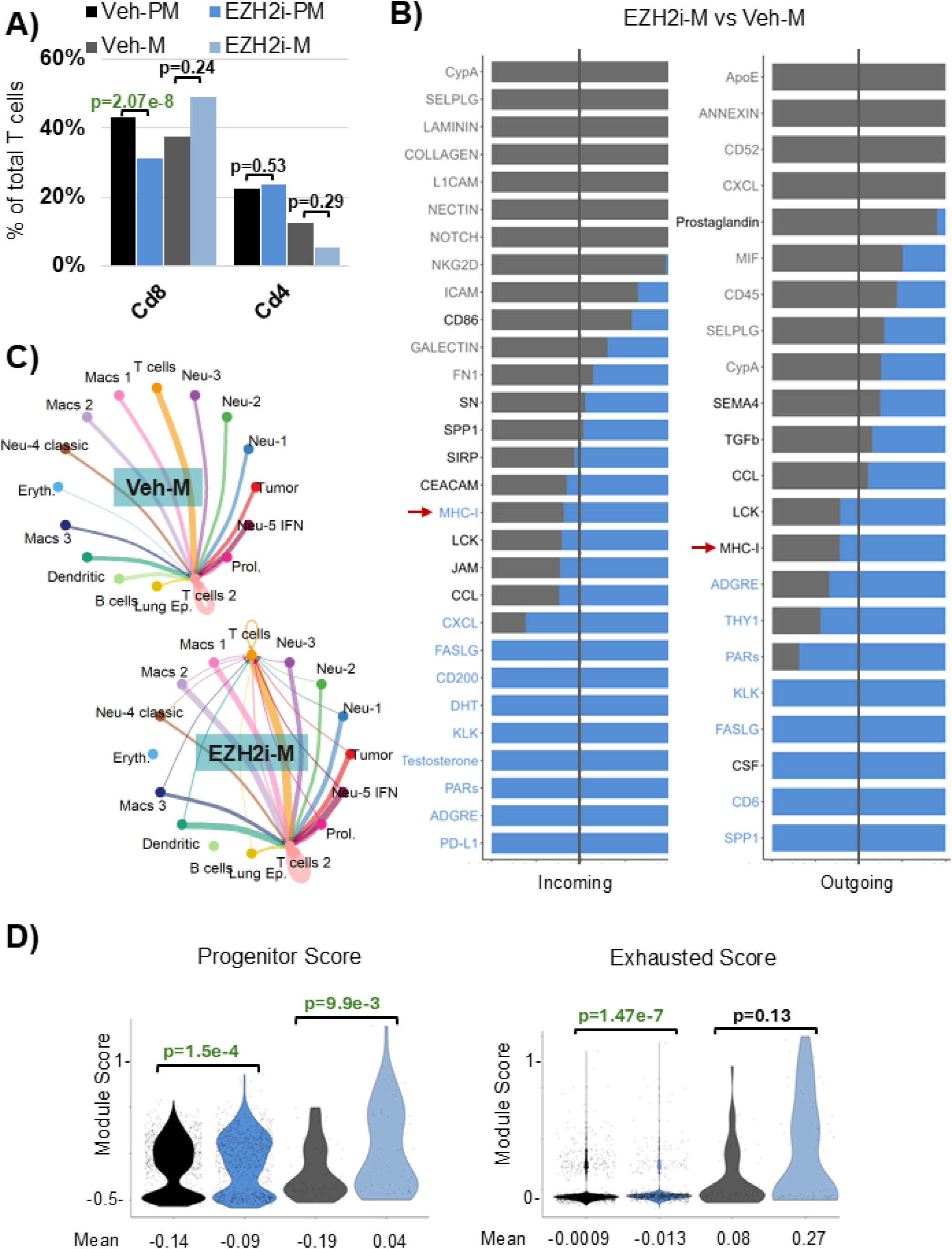
Endogenous murine TIL alterations. **A)** Proportions of Cd8+ and Cd4+ T cells. P-values reported are raw via Fisher’s exact test in R. **B)** Conserved and context-specific relative information flow alterations for highly significant (p<0.01) pathways incoming and outgoing from eTILs. Pathways with gray text are enriched in Veh-M, blue is enriched in EZH2i-M, and black indicates an enriched pathway in both groups. Red arrows are markers for pathways highlighted in remaining sub-figures. **C)** Circle plots of MHC-I signaling in Veh-M and EZH2i-M. Bar color corresponds with sender and size is proportional to the signal strength. **D)** Violin plots of module scores comparing progenitor (*Tcf7, Pdcd1, Havcr2*) and exhausted phenotypes (*Tox, Tox2, Pdcd1, Tigit*). P-values reported are raw values using the Mann–Whitney U test in R. Abbreviations: Neu – neutrophil; Macs – macrophage; Prol. – proliferating; Lung Ep – lung epithelia

CellChat revealed changes in eTILs from EZH2i-M that are indicative of an enhanced response, while also exhibiting modulatory effects. First, heightened antigen responsiveness and cytotoxic capacity due to immune synapse formation are seen with increased (or activated) incoming signals from LCK, MHC-I, FASLG, CEACAM, and CD6 (Figure-4B). Notice in Figure-4C that we now have MHC-I signaling from the whole TME to both T cell groups compared to vehicle which only communicates to Tcells2. Conversely, reduction (or elimination) of signals from LAMININ, COLLAGEN, FN1, SELPLG, and ICAM suggests extracellular matrix remodeling and a decrease in tissue retention. That is, eTILs may have more motility and access within the TME. Further, loss of NOTCH and NKG2D paired with decreased GALECTIN could indicate less terminal differentiation and exhaustion(^30^). Intriguingly, we see the potential emergence of compensatory mechanisms because there are clearly decreased immune suppression pathways like TGFb, Prostaglandin, CD52, and CypA, while simultaneously upregulating PD-L1 and CD200. This additionally supports the efforts of many ongoing trials combining EZH2i with ICI therapy. Lastly, outgoing signals such as FASLG, CSF, CD6, THY1, and SPP1 indicate a shift to activated and effector-like states, with cross-talk to myeloid compartments.

We next utilized modularity scores in Seurat to examine changes in phenotypes of eTILs - using Cell Signaling Technology’s established *Mouse Immune Cell Marker Guide*(^31^). A table of each module is available in **Supp. Table 1**. First, EZH2i significantly increases progenitor-like features in both treatment groups (PM: p=1.5e-4, Cliff’s δ = 0.096; M: p=9.99e-3, Cliff’s δ = 0.293), while exhaustion is significantly decreased in PM (p=1.47e-7, Cliff’s δ = -0.133) but modestly increased in M (p=0.13, Cliff’s δ = 0.17) – Figure-4D. We show additional eTIL phenotypes in **Supp. Figure-5** (effect sizes reported on GitHub), in short, EZH2i-PM contained significantly higher proportions of naïve eTILs (p=0.001), whereas EZH2i-M showed a non-significant reduction relative to vehicle (p=0.75). Activation followed a similar pattern: EZH2i-PM TILs were more activated than Veh-PM (p=0.042), while EZH2i-M were less activated than Veh-M (p=0.036). Cytotoxicity showed the inverse trend, with EZH2i-M exhibiting slightly greater cytotoxic potential than Veh-M (p=0.6) and EZH2i-PM displaying a mild reduction compared to Veh-PM (p=0.61). Both treatment groups had increased proportions of central memory eTILs compared to vehicle, though differences were not significant (PM: p=0.112; M: p=0.235). EZH2i-M contained significantly more effector-memory cells than Veh-M (p=0.0004), whereas EZH2i-PM contained significantly fewer than Veh-PM (p=0.0002). A similar trend is also seen within the effector populations (PM: p=0.006; M: p=0.123). This is presumably due to less prolonged effector activity in the premalignant TME.

Together, these data indicate that EZH2 inhibition drives distinct eTIL phenotypic programs depending on disease context. In the premalignant microenvironment, EZH2i promotes progenitor-like and naïve T-cell states with reduced exhaustion and effector differentiation, suggesting enhanced T-cell renewal potential. In contrast, within established malignancy, while EZH2i still promotes progenitor features, it favors more differentiated effector-memory and cytotoxic phenotypes, presumably reflecting increased antigenic stimulation and adaptation to a more suppressive TME. These findings highlight the context-dependent role of EZH2 in regulating T-cell differentiation and functional persistence across tumor progression stages.

## Human TIL exposure to transient Valemetostat

Given our *in silico* results from murine models, we moved to *in vitro* human NSCLC samples (Figure-5A(^32^)). Samples from two patients were collected and TILs were expanded using standard protocols(^33^). Assays were completed with at least three fragments per patient, and each fragment was considered a unique culture because each fragment has a unique TME. For example, in Figure-5B we show cell populations of H&E-stained tissue slides with the HALO® nuclear phenotyping algorithm. TILs expanded from a region similar to Figure-5B**.1** would likely be less robust than those expanded from Figure-5B**.5**. Therefore, each fragment is indeed its own replicate. Similar to prior work(^20^), we exposed TIL products to 300nM of valemetostat (Val) or vehicle (Veh) for 4 days prior to harvest to induce a transient reduction in H3K27me3.

**Fig. 5:**
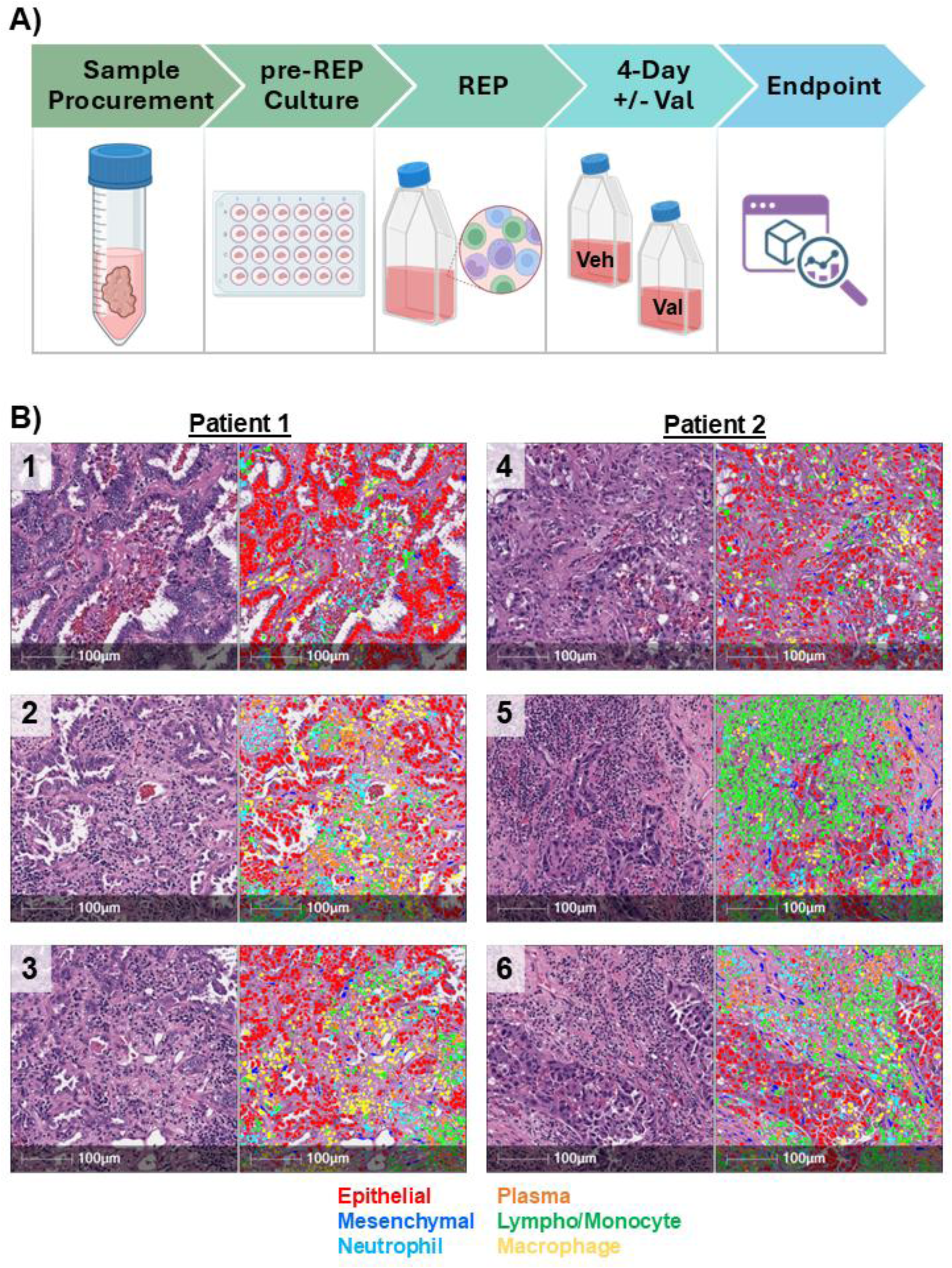
Human lung cancer TILs. **(A)** Schematic of workflow and design. **(B)** H&E and HALO nuclear phenotyper images showing cells within human NSCLC tumors.

Flow cytometry in Figure-6A&B revealed that Val treatment significantly shifted the CD8:CD4 ratio (0.7 vs 1.3; p=0.003). Val-treated cultures also exhibited a more effector memory phenotype (p=2.1e-4) with a corresponding decrease in central memory (p=8.4e-4). Proliferation, measured by Ki-67, was slowed in the Val group at harvest (p=1.9e-5), along with activation (Ki-67+/CD25+/CD69+, p=0.04) while CTLA-4 expression slightly increased (p=0.004). In contrast, other key phenotypic markers such as CD25, CD69, TCF1, and PD-1 were not significantly altered. Further gating strategies are shown in **Supp. Figure-6**.

**Fig. 6:**
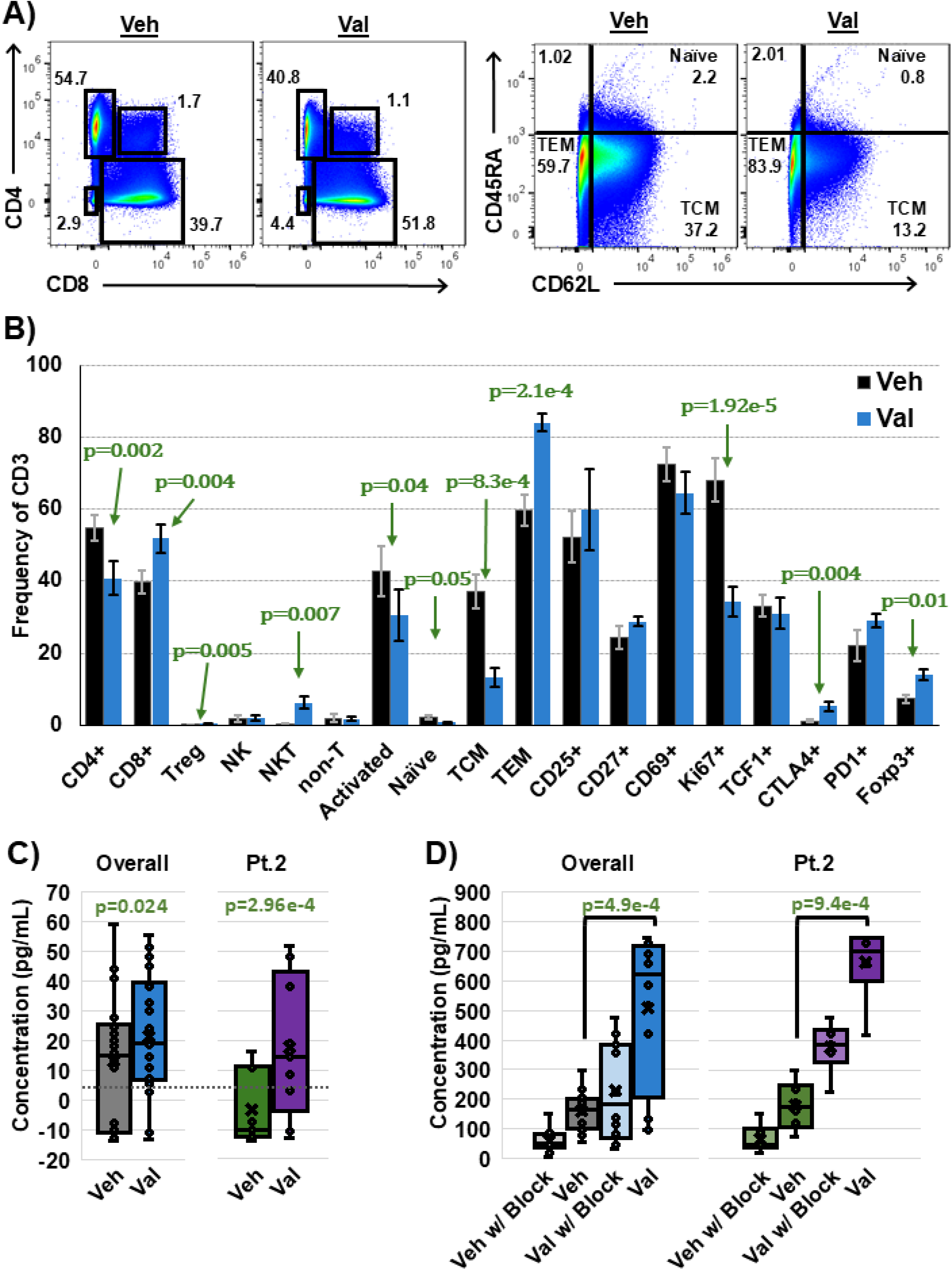
Phenotyping and reactivity of human lung cancer TILs. **(A)** Gating strategy for T cell subtypes and memory phenotype shown on concatenated respective treatment samples (Veh and Val). **(B)** Frequencies of total CD3, with P-values reported from paired two-tailed t-test. Error bars are SEM. **(C)** Allogeneic co-culture from two patients (pt. 2 shown) and 3 fragments among +/- Val (no MHC-block). P values are from a paired one-tailed t-test **(D)** Autologous co-culture from two patients (pt. 2 shown) and 3 fragments among +/- Val and +/- MHC-I block (W6/32). P values are from a paired one-tailed t-test. Treg: CD25+/Foxp3+/CTLA4+, NK: CD3-/CD56+, NKT: CD3+/CD56+, non-T: CD3-/CD56-, Activated: Ki-67+/CD25+/CD69+, Naïve: CD45RA+/CD62L+, TCM: CD45RA-/CD62L+, TEM: CD45RA-/CD62L-

Despite reduced proliferation at harvest, Val-treated TILs demonstrated greater functional reactivity upon tumor restimulation (Figure-6C&D), consistent with their increased proportion of effector memory cells. Due to limited autologous tumor material, we performed an allogeneic reactivity assay (Figure-6C) that showed enhanced IFNγ production following Val treatment quantified by IFNγ ELISA. Although overall magnitude of response was lower compared to the autologous assay (Figure-6D), likely due to MHC-restriction, the increase remained significant across patients (p=0.024) and was strongest in patient 2 (p=2.96e-4). Autologous reactivity was also significantly increased following Val treatment (p=4.9e-4), with patient 2 exhibiting the most robust response (p=9.4e-4). These results demonstrate that Val treatment enhances TIL functional reactivity following both tumor re-exposure and with allogeneic challenge.

## Discussion

In this study, we used single-cell resolution to gain mechanistic insights into how EZH2 inhibition alters the TME of NSCLC to become more immunogenic. We first demonstrated that cell-cell signaling was shifted to favor anti-tumor immune populations, where pathways such as MHC-I, FASLG, LCK, and CEACAM were more enriched and active in treatment groups. We also showed downregulation of fibrosis markers in COLLAGEN and FN1. Even though we inhibited EZH2 methyltransferase (H3kme3) activity, we demonstrated that *Ezh2* was still active. This is important because EZH2 has been shown to be essential for T cell development, where knockout leads to terminal differentiation of T cells. Concerning endogenous TIL populations, there were increases in CD8+ and decreases in helper CD4s. Pathway alterations revealed incoming and outgoing signaling associated with an improved immune response, and eTILs exhibited a more progenitor-like, less exhausted phenotype. These findings suggest that although EZH2 inhibition confers transient benefits, prolonged exposure may ultimately drive eTILs toward terminal differentiation or exhaustion.

We also expanded human NSCLC TILs under standard production protocols as a proof-of-concept and observed a pronounced shift toward an effector memory (T_EM_) phenotype with a corresponding reduction in central memory (T_CM_) following Val exposure. Concurrently, we saw the Val-TILs were much more reactive. Although T_CM_s are often considered more favorable for adoptive transfer due to superior persistence, lymphoid homing, and proliferative capacity, the importance of these traits may differ for lung-directed TIL therapy. Infused TILs rapidly localize in the lungs by nature of the pulmonary system, and reduced reliance on lymph node trafficking could potentially make a TEM-skewed product more advantageous if immediate effector function in the target organ is prioritized.

Nevertheless, T_EM_ enrichment can also reflect increased differentiation, which may limit long-term durability. Future studies assessing lung retention/migration and longer-term persistence will be important to determine whether this shift impacts therapeutic potential. While CD69 and TCF1 trended downward (non-significantly), further phenotyping is warranted to clarify whether Val alters progenitor-like versus terminal differentiation states. Upcoming panels will incorporate TIM-3 and CD39 to better resolve such subsets. Determining the optimal dosage and duration for maximizing TIL responsiveness remains an active area of investigation.

Notably, the EZH2 inhibitor used in the *in silico* study (GSK126) differs from the next generation of inhibitors moving forward in clinical trials (NCT05879484, NCT04102150, 24-LUN-140-DS-PMC). GSK126 failed in clinical trials due to its inability to inhibit tumor growth, a limitation that may partly stem from differences in drug design and availability compared to new inhibitors. GSK126 could only be used as an infusion, whereas Valmetostat (DS-3201) was specifically designed for oral delivery. Attempts to reformulate the compound for oral administration led to a substantial decrease in bioavailability, whereas Valmetostat (DS-3201) was specifically designed for oral delivery. They also differ in their mechanism of inhibition of EZH2. GSK126 and Valmetostat compete for S-adenosylmethionine (SAM) binding pocket that donates the methyl group allowing EZH2 to carry out its function, but Valemetostat is a dual inhibitor of EZH2/EZH1. Elucidating further mechanisms of Valmetostat could help to explain the variance in effectiveness between the first generation of EZH2 inhibitors and those currently in clinical trials.

While this study provides new systems-level insights into EZH2’s influence on tumor microenvironments (TMEs), several limitations warrant consideration. First, different resources rely on distinct databases that can lead to variable results across methods (e.g. results from SCENIC vs CollecTRI vs BITFAM). An additional limitation is the lack of an independent, publicly available scRNA-seq dataset that closely matches the experimental conditions used here. To our knowledge, no such dataset currently exists that would enable meaningful cross-dataset validation. Consequently, external validation using an independent single-cell cohort was not feasible and remains an important direction for future studies as comparable datasets become available. Second, underrepresentation of cell populations can reduce the resolution of downstream analyses, though key biological insights remain accessible. CellChat is constrained by its reliance on predefined ligand-receptor databases, which may not capture tissue-specific or context-dependent interactions. Inferred interactions also do not consider heterogeneity within cell groups (e.g. T cell sub-types), nor does CellChat account for potential post-translational modifications. SCENIC is inherently stochastic (inference with regression trees) and relies on pre-computed TF binding motif databases. This results in slight variability in identified regulons, targets, and weights between runs. It also cannot infer actions caused by post-translational activity. The SCENIC pipeline is also highly technical, complex, and not particularly user-friendly. Next, Seurat’s *AddModuleScore* function for calculating modularity scores is sensitive to gene set composition and size, with larger gene sets prone to inflated scores due to increased probability of capturing non-specific expression patterns. Finally, potential confounding factors include sex differences and the discrepancy in treatment duration between pre-malignant (two weeks) and malignant (four weeks) samples. To address these limitations, multiple complementary approaches were employed.

Future studies will focus on *in vitro* and *in vivo* impact of EZH2 inhibition on TIL production and efficacy, including migratory capacity, optimizing dosage, and calibrating timing for the best clinically translatable results. Other works have shown improvement for CAR-T and TCR cellular therapies, but little is known about appropriate dosage and timing with regard to the various phases of TIL production and infusion.

## Methods

### Animal work

*In vivo* murine studies were performed as previously described(^21^). Briefly, a mouse model of lung squamous cell carcinoma was generated by deleting both *Lkb1* and *Pten* floxed alleles through adeno-Cre inhalation. After approximately 40 weeks, a baseline MRI scan was performed and mice were assigned to treatment groups. The relevant ones for this study are vehicle (Captisol pH4.6) or GSK126 (300mg/kg) administered as intraperitoneal injections twice per week. Pre-malignant mice received treatment for two weeks while mice with lung tumors received treatment for four weeks. It is also important to note sex differences: Veh-PM (F), EZH2i-PM (F), Veh-M (F), and EZH2i-M (M). Thus, hormone shifts observed in CellChat through DHT and Testosterone are likely due to sex differences in the mice. All work was approved by the Institutional Animal Care and Use Committee (IACUC) at the University of Kentucky.

### Single-cell sequencing

Macro-dissected lung tumors from malignant mice, or whole lung tissue from premalignant mice, were collected, rapidly mechanically dissociated, and run over Miltenyi LS columns to obtain single cell suspension. Single-cell capture was performed with a 10x Genomics Chromium controller and the 10x Genomics Single Cell 3’ v3 Kit was used for reverse transcription. Libraries were then prepared and sequenced, and raw sequencing data were aligned to the mm10 mouse genome (v93) using the Cell Ranger v3.1 pipeline (10x Genomics). Downstream analysis was conducted using the Seurat (v3) R package(^34^). The workflow consisted of: (1) Quality control and preprocessing – removing genes detected in fewer than three cells, excluding low-quality cells with fewer than 100 detected genes, or those with over 7% mitochondrial transcript content. The filtered expression data were then log-normalized. (2) dataset integration and feature selection – log-normalized matrices from individual samples were merged, and highly variable genes were identified using the variance-stabilizing transformation (VST) method. The top 1,500 variable genes were used for integration. (3) Principal component analysis (PCA) was performed to reduce dimensionality, and cell clusters were identified using a shared nearest neighbor (SNN) graph-based clustering algorithm. (4) Cell identities were assigned by evaluating marker genes uniquely expressed in each cluster.

Differential expression analysis was performed using Seurat’s *FindMarkers* function with the Mann-Whitney U test to compare T cell gene expression between EZH2i-M vs Veh-M, EZH2i-PM vs Veh-PM, and EZH2i-M vs EZH2i-PM (**Supplementary**). Genes were considered differentially expressed if they had a log2FC>0.25, were expressed in at least 10% of cells in either group, and significance was determined by adjusted p-value < 0.05. Further, T cell clusters were compared using the Louvain algorithm and filtered to show -0.5<logFC>0.5, pct expression>0.1, and adjusted p-value<0.05.

Functional interpretation of genes, transcription factors (TFs), and pathways was primarily based on publicly available annotations. Unless otherwise stated, biological functions, pathway associations, and regulatory roles were retrieved from GeneCards (www.genecards.org) and NCBI Gene (www.ncbi.nlm.nih.gov/gene). These resources were used for general descriptive purposes and to provide standardized gene function summaries across datasets.

### CellChat

To infer and analyze intercellular communication networks, we used CellChat (v2.1.2), a computational tool that leverages scRNAseq data to model ligand-receptor signaling interactions between cell types using a curated database (KEGG and primary literature) (^22^). First, scRNAseq data was pre-processed as described above. The resulting expression matrices (split according to resistance group) from the Seurat object were used to create CellChat objects for each treatment arm according to the murine database. CellChat objects were then pre-processed to identify interactions and communication probabilities, including cell-cell and pathway-specific. Note that *interaction* refers to a specific ligand-receptor pair that mediates potential signaling between a sender cell type (expressing the ligand) and a receiver cell type (expressing the receptor). Each interaction is directional and context-dependent. Whereas communication *strength* is a quantitative measure reflecting the total signaling probability between two cell types, often summed over all interactions in a given signaling pathway. It integrates the expression levels of all relevant ligand-receptor pairs and provides a global metric of pathway activity using the law of mass action. All code were written in accordance with the author’s vignettes in Rstudio (v4.4.3) (https://github.com/sqjin/CellChat). Significant signaling pathways were identified based on permutation tests and statistical filtering (e.g., p-value < 0.05 unless otherwise stated).

### SCENIC

Transcriptional regulatory networks and assessment of transcription factor (TF) activity at the single-cell level was inferred using the Single-Cell rEgulatory Network Inference and Clustering (SCENIC) pipeline with the same top 1500 variable features retrieved through integration. Compared to alternative methods, SCENIC is advantageous because it combines expression analysis with motif enrichment to infer direct (versus correlative) TF-target relationships. In particular, we used the pySCENIC implementation (v0.12.1 - https://github.com/aertslab/pySCENIC) that is performed in three main steps using Spyder (v5.5.1, Python 3.12.7).

First, pySCENIC infers gene co-expression modules using GRNBoost2, a regression-based algorithm that identifies potential regulatory relationships between transcription factors and their target genes based on expression patterns across the dataset. This step generates a list of candidate TF-target interactions without assuming direct binding. Second, these preliminary regulons are refined (i.e. pruned) using cisTarget, which integrates cis-regulatory motif enrichment and genomic annotation data. Only targets with enriched motifs for the corresponding TF are retained, resulting in a high-confidence set of transcription factor–centered modules (also called regulons). Lastly, cells are differentiated and clustered according to regulon activity using AUCell. AUCell calculates the area under the cumulative recovery curve (AUC) for each regulon’s gene set in a cell’s ranked gene expression profile. This provides a quantitative estimate of regulon activity on a per-cell basis, enabling the identification of cell-type–specific transcriptional programs and dynamic regulatory changes across different cell states.

As with CellChat, treatment groups were analyzed separately and then compared, but with data converted from Seurat objects to loom format for compatibility with pySCENIC. The analysis utilized mouse-specific regulatory databases including ranking databases for cis-regulatory motif analysis (mm9-500bp-upstream, mm9-tss-centered-5kb, and mm9-tss-centered-10kb with 10-species conservation) and a comprehensive transcription factor list (allTFs_mm.txt) containing mouse TF annotations. The three main steps of the pipeline were implemented as described above and Regulon Specificity Scores (RSS) were calculated to identify cell type-specific transcription factors. We also calculated regulon similarities based on target gene overlap using Jaccard similarity metrics.

### Complementary methods

Several tools have been developed to infer cell-cell communication from scRNAseq data. However, these often rely on single ligand-receptor pairs, overlooking that many receptors form multi-subunit complexes. Methods such as CellPhoneDB(^35^) improve this by accounting for receptor subunit expression but still omit key cofactors including soluble modulators and co-receptors. Additionally, many existing tools lack curated pathway classifications, comprehensive network visualization, support for complex systems analysis, and the ability to capture communication across dynamic cell states. We employed the LIANA(^23^) (LIgand-receptor ANalysis frAmework) that provides a unified interface for inferring cell-cell communication from single-cell RNA sequencing data by integrating multiple ligand-receptor inference methods and databases. The framework decouples computational methods from their corresponding resources, enabling any combination of 7 different scoring algorithms with 16 distinct ligand-receptor databases, and generates consensus predictions across methods to reduce individual method bias.

BITFAM(^25^) (Bayesian Inference Transcription Factor Activity Model) is a Bayesian hierarchical framework that infers TF activities in individual cells by integrating scRNAseq data with prior knowledge from ChIP-seq transcription factor binding experiments. The method decomposes log-normalized expression data into matrices representing transcription factor activities and target gene weights using variational inference, where ChIP-seq data provides regulatory prior knowledge through binary matrices indicating predicted target genes for each transcription factor. BITFAM operates on the biological principle that differences in single-cell gene expression profiles reflect distinct underlying transcription factor activity states, enabling identification of regulatory mechanisms and cell clustering based on inferred transcription factor activities rather than gene expression alone.

Lastly, we used CollecTRI(^26^) (Collection of Transcriptional Regulatory Interactions) which is a comprehensive meta-resource containing signed transcription factor-target gene interactions compiled from 12 different curated databases, providing regulatory relationships for 1,186 transcription factors with mode-of-regulation annotations (activation or inhibition). The resource integrates information from multiple knowledge bases using a systematic workflow to assign regulatory signs to transcription factor-gene interactions, resulting in enhanced coverage and performance in identifying perturbed transcription factors compared to individual databases.

### Module scores

Module scores were calculated using the *AddModuleScore()* function in the Seurat R package (v 5.3.0). This method quantifies the relative expression of a predefined gene set within individual cells, facilitating the assessment of pathway activity at the single-cell level. First, we defined our gene set of interest according to phenotypic markers (e.g. *Sell+/Cd44-*). To account for both positive and negative markers, combined scores were computed by creating a separate module for negative markers and then subtracting this module score from the positive marker module score. For each cell, the average expression levels of these genes were computed to obtain a module gene expression score. Considering technical variability, including differences in sequencing depth and baseline expression, Seurat generates a control gene set. This background set consists of randomly selected genes matched to the module genes by overall expression level. The final module score for each cell was then calculated as the difference between the mean expression of the module genes and the mean expression of the matched control genes.

### GRN inference and analysis

Regulatory networks were inferred from scRNAseq data using GRNboost2(^36^), which employs gradient-boosting machines to model expression of genes based on candidate TFs. For every target gene, it constructs a regression model using shallow decision trees, where each tree is trained on a random subset of the data (90%) and evaluated on the remaining 10%. New trees are added iteratively until the average improvement of prediction falls below zero. Importance scores of TFs in predicting target gene expression are aggregated across all models to produce a final ranked regulatory network.

Once a GRN was created, analysis was performed using NetworkX (v3.4.2) and Pyvis (v0.3.2) in Python. The top 500 TF-target interactions ranked by importance score were used to construct directed and weighted (but not signed) networks. Nodes were classified as transcription factors (TFs), target genes, or dual-role genes (acting as both TF and target), with network topology analyzed through standard graph theory metrics including degree distribution, connectivity patterns, and component structure. Weakly and strongly connected components were identified to assess network fragmentation, while centrality measures (betweenness and closeness centrality) were calculated to identify key regulatory hubs and bridge nodes. Edge weight statistics characterized the distribution of regulatory interaction strengths, and network motifs including self-loops (auto-regulation) and mutual regulation pairs were quantified. Master regulators were identified as nodes with the highest out-degree (number of target genes regulated), while the most regulated genes were determined by in-degree (number of incoming regulatory connections). Interactive network visualizations were generated with Pyvis, incorporating physics-based layout algorithms and node/edge styling based on functional classifications and interaction strengths.

Phenotype control theory using feedback vertex set (FVS)(^37^) was also used. As mentioned in the **Supplementary** file, the inferred GRN has no inherent update functions, and we must rely on topological structures to find targets. In FVS control, by pinning a subset of nodes that intersect every cycle in the network, we disrupt all feedbacks, making the resulting network admit a single steady state. In other words, a FVS of a graph is a minimal set of nodes whose removal leaves the graph without cycles(^29^).

### Trajectories and pseudotime

Trajectory analysis was performed using Monocle3(v1.3.7) to infer differentiation velocities within T cell populations. T cell subsets were extracted from the integrated Seurat object, merged according to vehicle or treatment (because overall counts were insufficient to analyze PM/M separately), and converted to a *cell_data_set* object. Dimensionality reduction was performed using principal component analysis followed by UMAP embedding. Cells were clustered using the Louvain algorithm and assigned to partitions. Trajectory graphs were learned using the SimplePPT algorithm, and cells were ordered along pseudotime after manual selection of root nodes corresponding to naive/early T cell states based on *Tcf7*-high regions. Differential gene expression along the trajectory was assessed using Moran’s I test to identify genes with significant expression changes as a function of pseudotime. All analyses were performed separately for each treatment group to enable comparative trajectory analysis.

### TIL culture

All cultures were maintained according to University of Kentucky (UK) biosafety guidelines, as approved by the Institutional Review Board (#79432). Informed consent statement: Written informed consents were obtained and approved at the time of sample collection by UK’s Biospecimen Procurement and Translational Pathology Shared Resource Facility (BPTP SRF).

For the initial TIL expansion phase (pre-REP), TILs from fresh NSCLC tumor fragments (2-8mm^3^) were cultured in 24-well plates in TIL Complete Media (CM) with 6000 IU/mL IL-2 (Teceleukin, BRB Preclinical Biologics Repository, Frederick National Laboratory for Cancer Research) for 2-4 weeks. Media used for TIL expansion consisted of RPMI 1640 with L-glutamine, 10% heat-inactivated human AB serum, 10 mM HEPES, 100 IU/mL penicillin, 100 ug/mL streptomycin, 55 uM 2-mercaptoethanol, and 62.5 mg/mL amphotericin-B. Half of the media was replaced at least every 3-4 days, and the wells were sub-cultured when 80% confluent. For the rapid expansion protocol (REP), TILs were stimulated with 30 ng/mL anti-human CD3 antibody (Clone OKT3, Frederick National Laboratory for Cancer Research) in the presence of 3000 IU/mL of IL-2 and allogeneic irradiated PBMCs (50 Gy at 200x) from at least 10 different donors in 50/50 TIL-CM and AIM-V (Thermo #12055-091). TILs were split and media was replaced as needed. Four days prior to harvest, flasks were split equally into groups of vehicle (Veh) and Valemetostat (Val; MedChemExpress # HY-109108), where respective flasks received DMSO or 300nM Val. Cells were harvested on days 14-16.

### Histology and Halo

Tissues were embedded in paraffin and sectioned at the Markey Cancer Center BPTP SRF. Hematoxylin and eosin (H&E)-stained slides were scanned with an Aperio slide scanner, then were used for HALO® nuclear phenotype image analysis (Indica Labs - previously trained to recognize cell types(^21^)) at 20X magnification.

### Reactivity

Expanded TILs (post-REP) were co-cultured with autologous or allogeneic tumor single-cell suspensions for 18-24 hours at a 1:1 ratio in 96-well plates (VWR #10062-900). Tumor reactivity was quantified by measuring IFNγ levels in culture supernatants using ELISA (BioLegend, #430104). Allogeneic co-cultures were performed with or without valemetostat-treated TILs with n=3 replicates per sample per cell line.Autologous co-cultures for each patient were performed in vehicle-treated conditions (n=6) and Val-treated conditions (n=12; technical replicates were pooled and averaged for statistical analysis). For allogeneic co-cultures, reactivity was defined as IFNγ secretion above 4 pg/mL, corresponding to the lowest detectable level reported by the manufacturer; MHC blocking was not assessed due to the inherent MHC restriction in allogeneic settings. Allogeneic co-cultures were performed using three tumor cell lines (A549, PC9, and osimertinib-resistant PC9), kindly provided by Christian Gosser and Dave-Preston Esoe. Reactivity in autologous co-cultures was defined as IFNγ secretion exceeding 50 pg/mL and demonstrating at least 20% inhibition in the presence of an HLA-A,B,C blocking antibody (BioLegend, clone W6/32, #311402).

### Flow

Samples were analyzed using a CytekAurora spectral flow cytometer (BD Biosciences) and FlowJo v10.10 software. Samples were washed and acquired with at least 5x10^5^ events per sample. Staining and acquisition were completed by University of Kentucky’s Flow Cytometry and Immune Monitoring Core Facility. In short, cells were stained with antibodies at 1:100 in FACS buffer containing 0.5% FBS, 0.1% NaN3, and 0.5mM EDTA. Fc receptors were blocked with human IgG Fc blocking antibody (TruStain FcX, Biolegend). Surface staining was performed with CD69-BUV737 (FN50, BD Biosciences), CD45RA-eFluor450 (H100, Cytek), CD3-BV570 (UCHT1, Biolegend), CD62L-BV605 (DREG, Biolegend), CD28-BV650 (28.2, Biolegend), CTLA-4-BV711 (L3D10, Biolegend), CD56-BV750 (5.1H11, Biolegend), PD-1-BV785 (EH12.2H7, Biolegend), CD8-eFluor BG532 (SK1, Cytek), CD4-cFluro YG1 (SK3, Cytek), CD25-PE-Fire 700 (M-A251, Biolegend), anti-human IgG4-AF647 (H6025, SouthernBiotech), and viability dye (Zombie NIR, Biolegend). Following surface staining, cells were fixed and permeabilized with FOXP3 fixation/permeabilization buffer (eBioscience #00-5521-00), then stained with intracellular markers TCF1-RB705 (C63D9, BD Biosciences), Ki-67-RB744 (B56, BD Biosciences), and FOXP3-R718 (236A/E7, BD Biosciences).

### Statistics and reproducibility

Statistical analyses were performed using Python, R, or Excel. Each primary inference method used in this study incorporates internal statistical procedures, including multiple-testing adjustments and thresholding; details are available in the corresponding publications (and all linked in GitHub). For pre-planned independent pairwise comparisons, we report raw p-values, as multiple comparisons across groups did not occur (e.g., Figure-4). Cell-population proportions between paired treatment groups were compared using Fisher’s exact test. Comparisons of module scores and gene regulatory network (GRN) importance between paired treatment groups were performed using the Mann–Whitney U test. Results with Benjamini–Hochberg adjustments are not reported because cross-group multiple comparisons were not performed, but adjusted p-values are available upon request. Normality was assessed using the Shapiro–Wilk test, and effect sizes were calculated using Cliff’s δ. TIL phenotyping comparisons used a student’s paired two-tailed t-test, and reactivity was assessed using a student’s paired one-tailed t-test due to the directional hypothesis of increased reactivity.

## Data availability

All data presented in this article are available from the corresponding author upon reasonable request. The sequencing data are available at NCBI Gene Expression Omnibus under the super-series GSE233665, and full method-specific results are available at (https://github.com/drplaugher/EZH2-NSCLC-scRNAseq-TILs).

## Code availability

All code used in this article are available from the corresponding author upon reasonable request. The scripts build upon publicly available vignettes provided by the respective original authors in their published repositories. Modifications and additions specific to this work are available on the GitHub repository, along with instructions for reproducing the results. Proper attribution to the original sources is provided within the repository.

## Acknowledgements

This work was supported in part by University of Kentucky’s Markey Cancer Center Shared Resource Facilities (P30-CA177558): Biostatistics & Bioinformatics, Biospecimen Procurement & Translational Pathology, Flow Cytometry & Immune Monitoring. Services in support of the research project were provided by the VCU Massey Comprehensive Cancer Center Bioinformatics Shared Resource. Massey is supported, in part, with funding from NIH-NCI Cancer Center Support Grant P30 CA016059.

## Funding

This work was supported in part by NCI K99CA303792 (DRP), NCI T32-CA165990 (DRP, CMG), T32 ES007266-30 (TD), National Center for Advancing Translational Sciences - NIH UL1TR001998 (DRP, CFB), R01-CA237643 (CFB), and R01-HL170193 (CFB). The content is solely the responsibility of the authors and does not necessarily represent the official views of the NIH.

## Author Contributions

Conceptualization: DRP, CFB; Data Curation: DRP; Data acquisition and processing: DRP, TD, JL, JinzeL, XQ, YL, CMG, CFB; Formal analysis: DRP, AC, CMG, CFB; Bioinformatics and biostatistics: DRP, JL, JinzeL, XQ; Writing – original draft: DRP, CFB, AC; Writing – review and editing: DRP, CFB, AC, CMG, DPE

## Ethics Declarations

TD was employed by Caris Life Sciences after completion of the work described in this manuscript. Caris Life Sciences had no role in the study design, data collection, analysis, interpretation, or manuscript preparation. The remaining authors declare no competing interests.

## Supplementary information

See the **Supplementary File** and GitHub repository.

## SUPPLEMENTARY MATERIALS

Daniel R. Plaugher et. al.

## Supplementary Figures 1-6

The computational workflow, supplementary analyses, and datasets supporting this study are publicly available in a GitHub repository (https://github.com/drplaugher/EZH2-NSCLC-scRNAseq-TILs). The repository contains analysis scripts written in R and Python for cell-cell communication analysis using CellChat and LIANA, transcription factor regulatory network inference using pySCENIC, BITFAM, and CollecTRI, and trajectory analysis using Monocle3. Supplementary figures documenting cell signaling patterns, transcription factor activities, statistical summaries, and additional complementary analyses are organized into dedicated directories. Raw sequencing data have been deposited in the Gene Expression Omnibus (GEO) under accession number GSE233665, while large pre-processed datasets and derived results are available through Zenodo (DOI: 10.5281/zenodo.15579657). All code is released under the MIT License and datasets are available under CC BY 4.0, ensuring full reproducibility and transparency of the analytical methods employed in this study.

**Supplemental Fig. 1:**
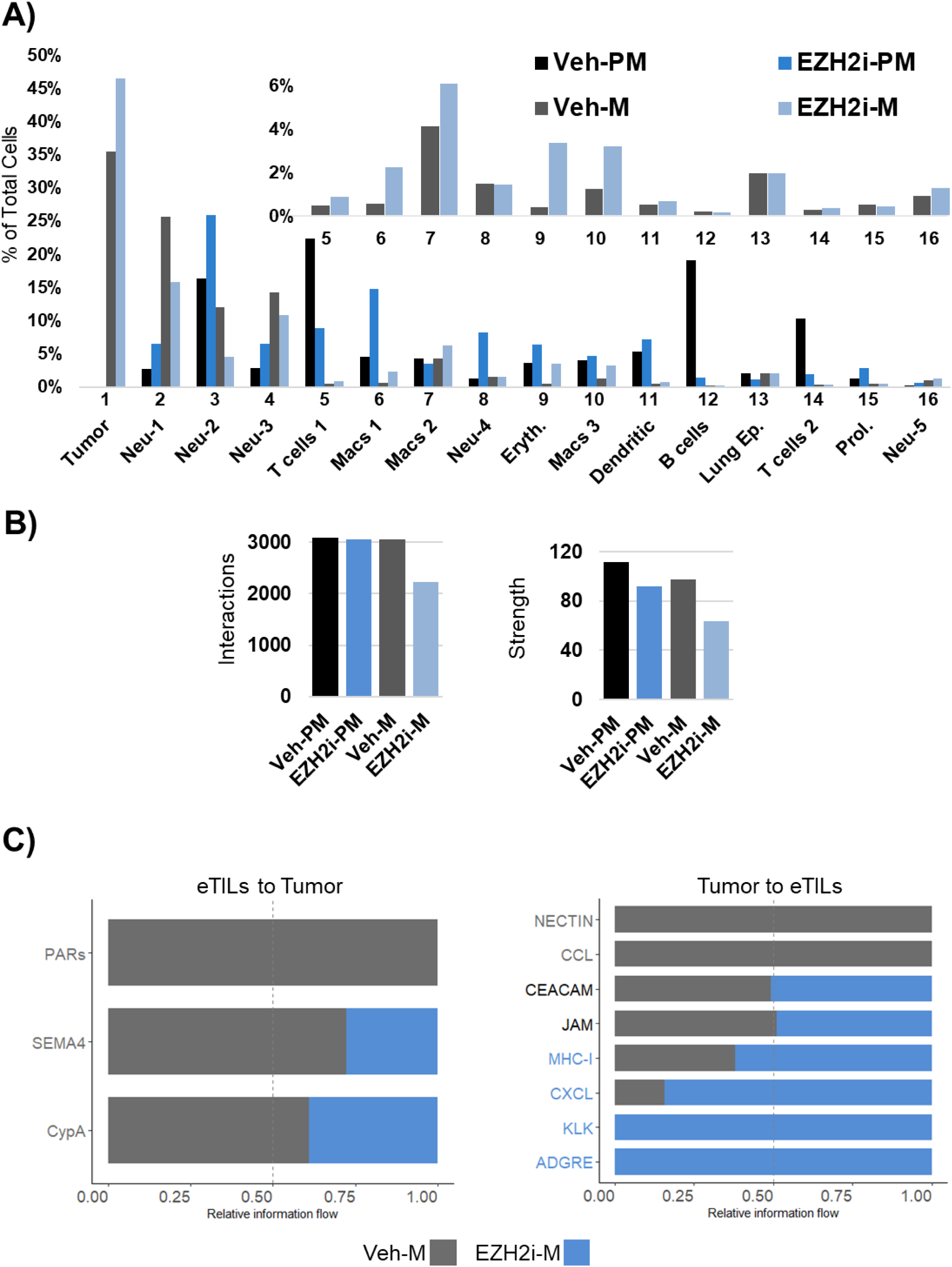
Differential Signaling between Tumor and T cells. **A)** Percentage of annotated cells per treatment group graphed for all populations, zoomed subgraph only shows Veh-M and EZH2i-M. **B)** Total number of interactions and interaction strengths of the inferred cell-cell communication networks between conditions. **C)** Relative information flow for significant pathways (p<0.05) between T cells and Tumor. Pathways colored gray are enriched in Veh-M, blue is enriched in EZH2i-M, and text colored black indicates an enriched pathway in both groups.

### EZH2 inhibition alters transcription factor activity

Overall, each treatment group has distinct activity, shown with a max-normalized distribution (‘TF signaling’ folder on GitHub). First, EZH2 inhibition does not appear to dramatically shift the number of active translational programs, but rather the factors that make up their regulons. Veh-PM has 46 regulons, an average target magnitude of 130.3 (largest is *Tfec* with 514 targets), and is led by *Mef2c*, *Ebf1*, and *Pax5* – favoring B cell development. Veh-M has 43 regulons, an average target magnitude of 142.1 (largest is *Irf8* with 656 targets), and is led by *Egr1*, *Foxb*, and *Atf3* – indicating stress response and differentiation. EZH2i-PM has 49 regulons, an average target magnitude of 122.9 (largest is *Irf8* with 603 targets), and is led by *Ddit3*, *Jdp2*, and *Atf3* – favoring stress response and apoptosis. Lastly, EZH2i-M has 47 regulons, an average target magnitude of 123.4 (largest is *Irf8* with 625 targets), and is led by *Elf5*, *Yod1*, and *Irf7* – indicating interferon response and inflammation.

Relative to each treatment arm, we see that EZH2i has less enriched TF activity compared to vehicle (Supp **FIG-2**). Examination of greatest changes in log2FC shows that EZH2-PM versus Veh-PM has decreases in *Pbx1* (−5.4), *Ebf1* (−4.9), and *Pax5* (−4.5), with increases in *Jdp2* (1.5), *Nr1h3* (1.4), and *Ddit3* (1.4). That is, we see a decrease in B cell differentiation factors and an increase in stress response, lipid metabolism, and apoptosis. EZH2-M versus Veh-M has decreases in *Fosb* (−5.8), *Nfia* (−2.7), and *Ebf1* (−2.1), with increases in *Cebpa* (4.4), *Maf* (1.9), and *Atf5* (1.6). These show a decrease in developmental factors with a shift in myeloid differentiation and stress response.

**Supplemental Fig. 2:**
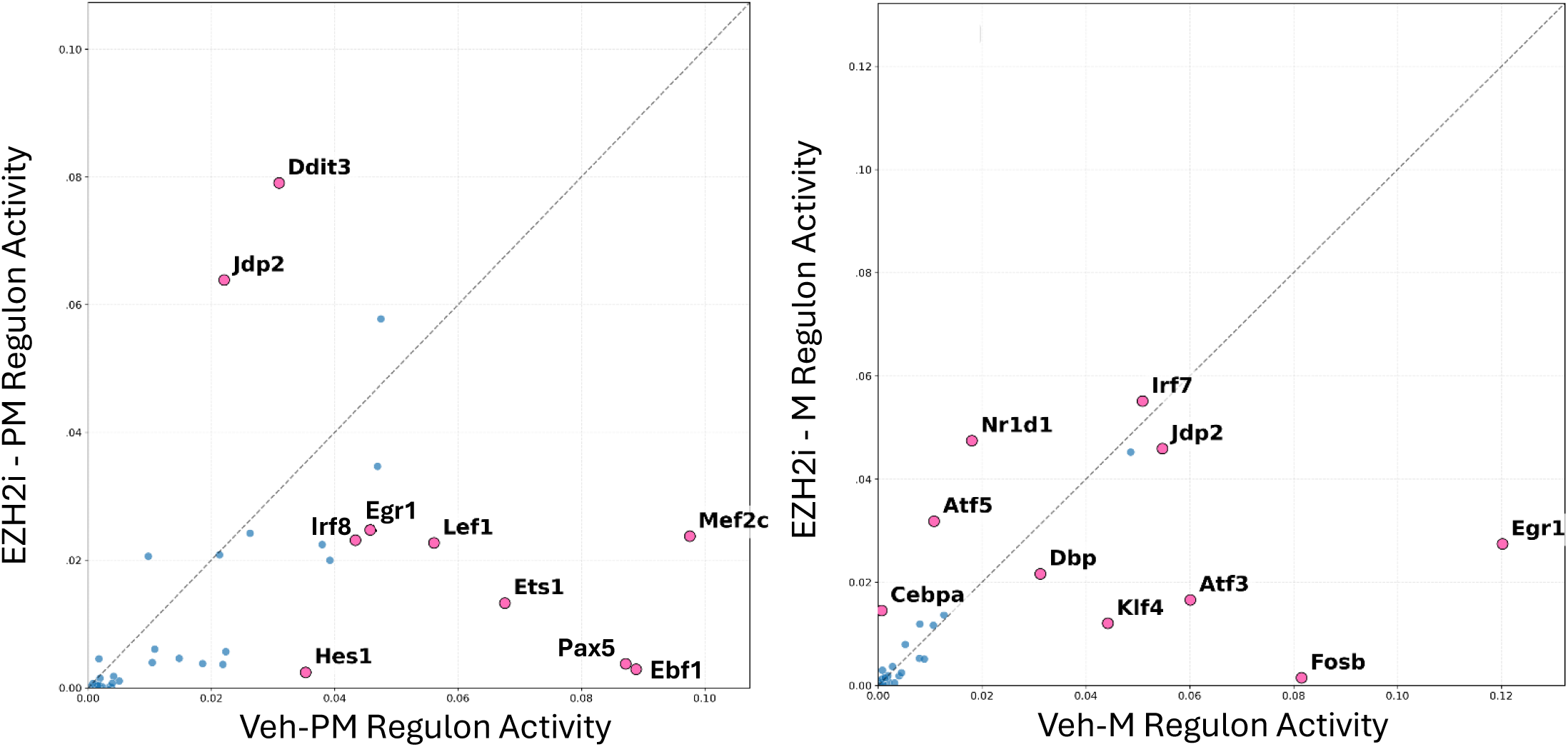
Scatter plots comparing shared TF activity between treatment (y) and vehicle (x) among pre-malignant and malignant conditions. TFs highlighted are the top 10 most differentially active

**Supplemental Fig. 3:**
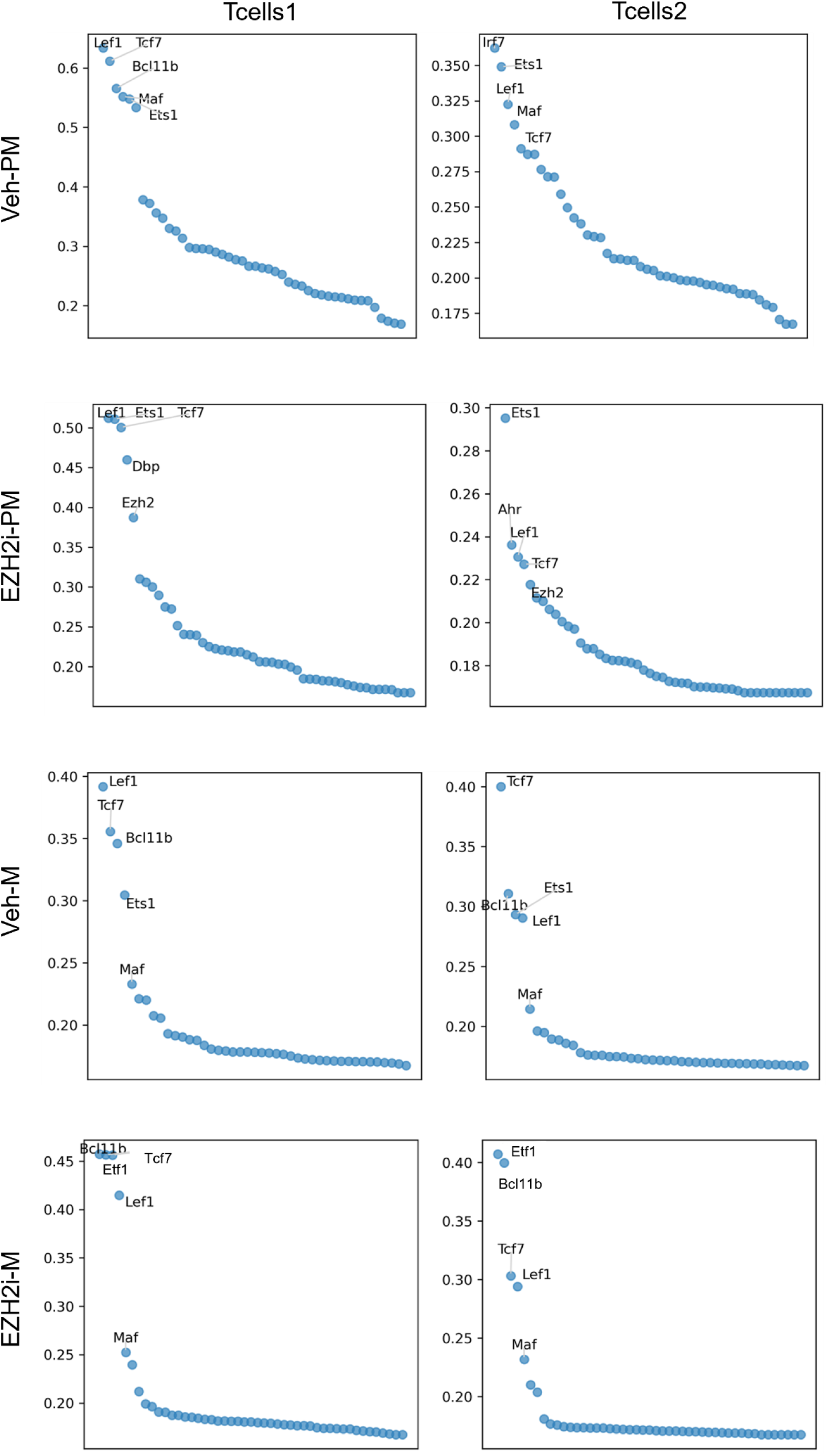
Regulon specificity scores for T cells among conditions. The RSS quantifies how

### Gene regulatory networks become more granular with EZH2 inhibition

As part of the pySCENIC pipeline, GRNboost2 generates directed and weighted TF-target estimates that we used to build gene regulatory networks (GRNs). We have taken these signals and converted them into interactive GRNs of the top 500 communications using Pyvis, then performed network analysis for connectivity, connected components (weak and strong), structure, centrality, and motifs. Since these networks are too large to display within the article, we have linked them to the GitHub repository. There we indicate target genes (green), transcription factors (blue), and dual-role actors (purple). Comparing cohorts with malignant TMEs, we first see that EZH2i makes the GRN more fragmented, as the 500 interactions are spread between 477 unique genes (53 TF only, 398 target only, 26 dual-role) compared to 440 in Veh-M (49 TF only, 367 target only, 23 dual-role).

These interactions contain 38 components (largest 269, 7 multi-node strong components) and 30 (largest 303, 5 multi-node strong components) for treatment and vehicle, respectively. Importance scores have means of 7.9 for vehicle and 7.3 for treatment (p=0.07, Cliff’s δ = 0.067). Regulators with the highest out-degree between both cohorts remain similar, however, slight decreases in *Irf7* and *Arg1* are observed while having increases in *Irf8*, *Zeb2*, and *Bhlhe41*. This may indicate a decrease in immune suppression through polarization of myeloid populations and enhanced immune signaling. Further, EZH2 inhibition leads to the emergence of mutual regulation of *Lef1* and *Tcf7* – possibly indicating the reprogramming of immune cells into a more stem-like phenotype.

Exemplifying the complex nature of EZH2 communications, we again implemented GRNboost2 to generate interaction networks focused around the top 100 *Ezh2* interactions where edges are weighted by importance scores (interactive version on GitHub). Veh-M has an average importance score of 3.04 but EZH2i-M has an average of 1.94 (p = 4.66x10^-11^, Cliff’s δ = 0.54). Thus, EZH2 inhibition substantially weakens EZH2’s regulatory influence while rewiring its interaction network.

A useful feature of dynamical systems analysis is that elements of the GRN can be targeted to drive the system to a preferred state using *phenotype control theory* (^29^). Given that our GRNs are data-driven, they do not inherently contain functional equations (i.e. update rules) to use methods like computational algebra (^38^) or stable motifs (^39^). However, strongly connected components (SCC) allow us to infer potential targets with a feedback vertex set using strictly the topology of a network(^37^). Given the SCC of the Veh-M network (diseased state), targets such as *Arg1* (immune suppression) or *Hspa5* (tumor survival) arise as potential therapeutic vulnerabilities of the overall system. Arginase 1 (ARG1) suppresses immune response by catabolizing arginine, which is essential for T cell activation and proliferation(^40^). HSPA5 (heat shock 70kDa protein 5) is an endoplasmic reticulum (ER) chaperone that is highly expressed under ER stress and protects cancer cells from ferroptosis and apoptosis(^40^). Importantly, while these elements still exist in the treatment GRN, they are much less prevalent and influential.

**Supplemental Fig. 4:**
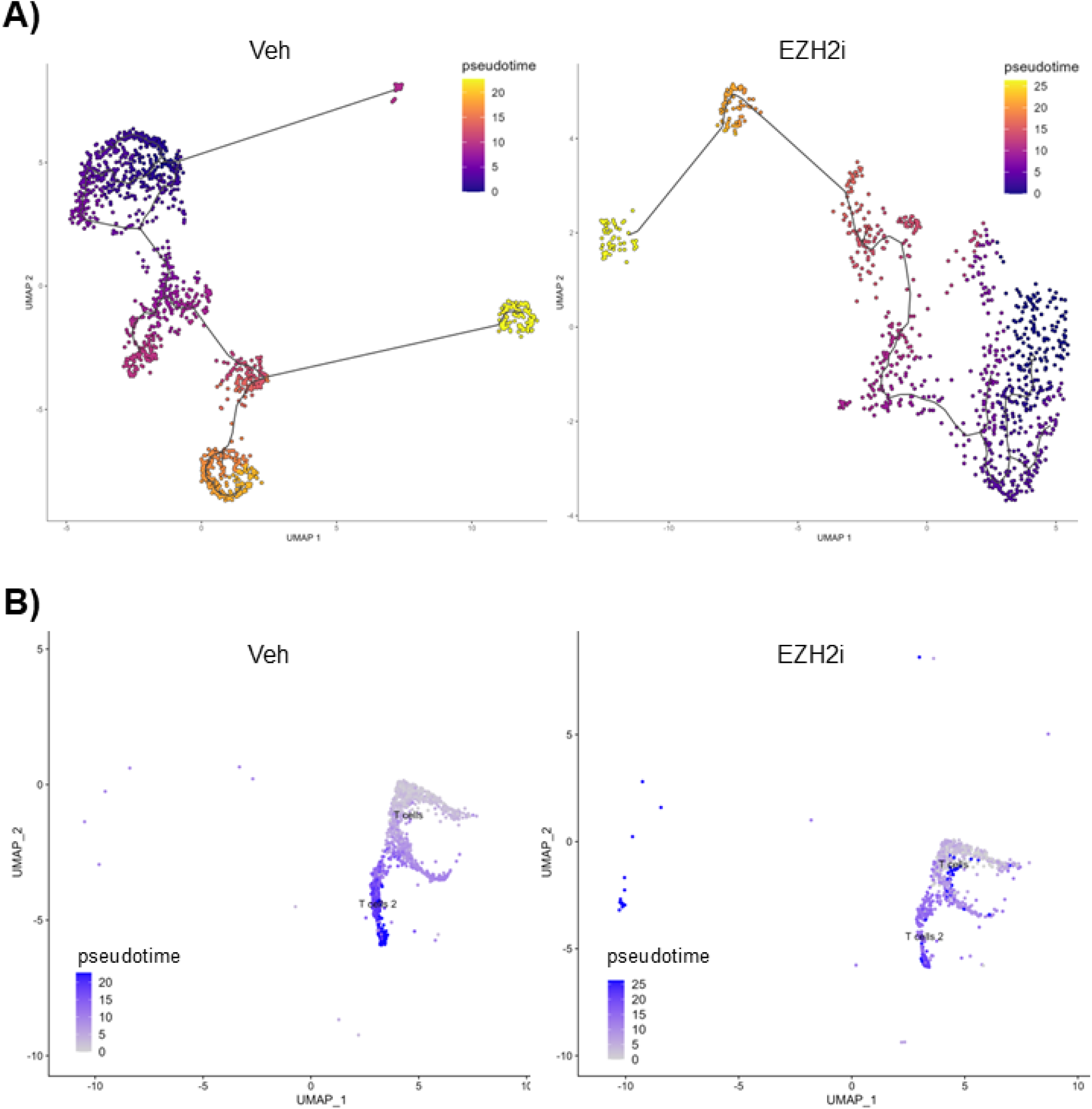
Trajectory analysis of T cell differentiation. **A)** Inferred clustering with trajectories and pseudotime, using Tcf7 to identify root nodes. Due to low counts, treatment conditions (PM/M) were merged – likewise with vehicle. **B)** Resulting pseudotime from (A) is overlayed with the original UMAP of T cell populations.

**Supplemental Table 1:** Markers used for modularity scores. Human orthologs are as follows - Cd69: CD69, Il2ra: CD25, Gzma: GZMA, Gzmb: GZMB, Prf1: PRF1, Sell: CD62L, Cd44: CD44, Il7r: CD127, Cd44: CD44, Il7r: CD127, Cd44: CD44, Sell: CD62L, Cd44: CD44, Sell: CD62L, Tcf7: TCF1, Pdcd1: PD-1, Havcr2: TIM-3, Tox: TOX, Tox2: TOX2, Pdcd1: PD-1, Tigit: TIGIT

| Phenotype | Positive/High | Negative/Low |
| --- | --- | --- |
| **Activated** | *Cd69, Il2ra* |  |
| **Cytotoxic** | *Gzma, Gzmb, Prf1* |  |
| **Naive** | *Sell* | *Cd44* |
| **Effector_mem** | *Il7r, Cd44* | *Sell* |
| **Central_mem** | *Il7r, Cd44, Sell* |  |
| **Effector** | *Cd44* | *Sell* |
| **Progenitor** | *Tcf7, Pdcd1, Havcr2* |  |
| **Exhausted** | *Tox, Tox2, Pdcd1, Tigit* |  |

**Supplemental Fig. 5:**
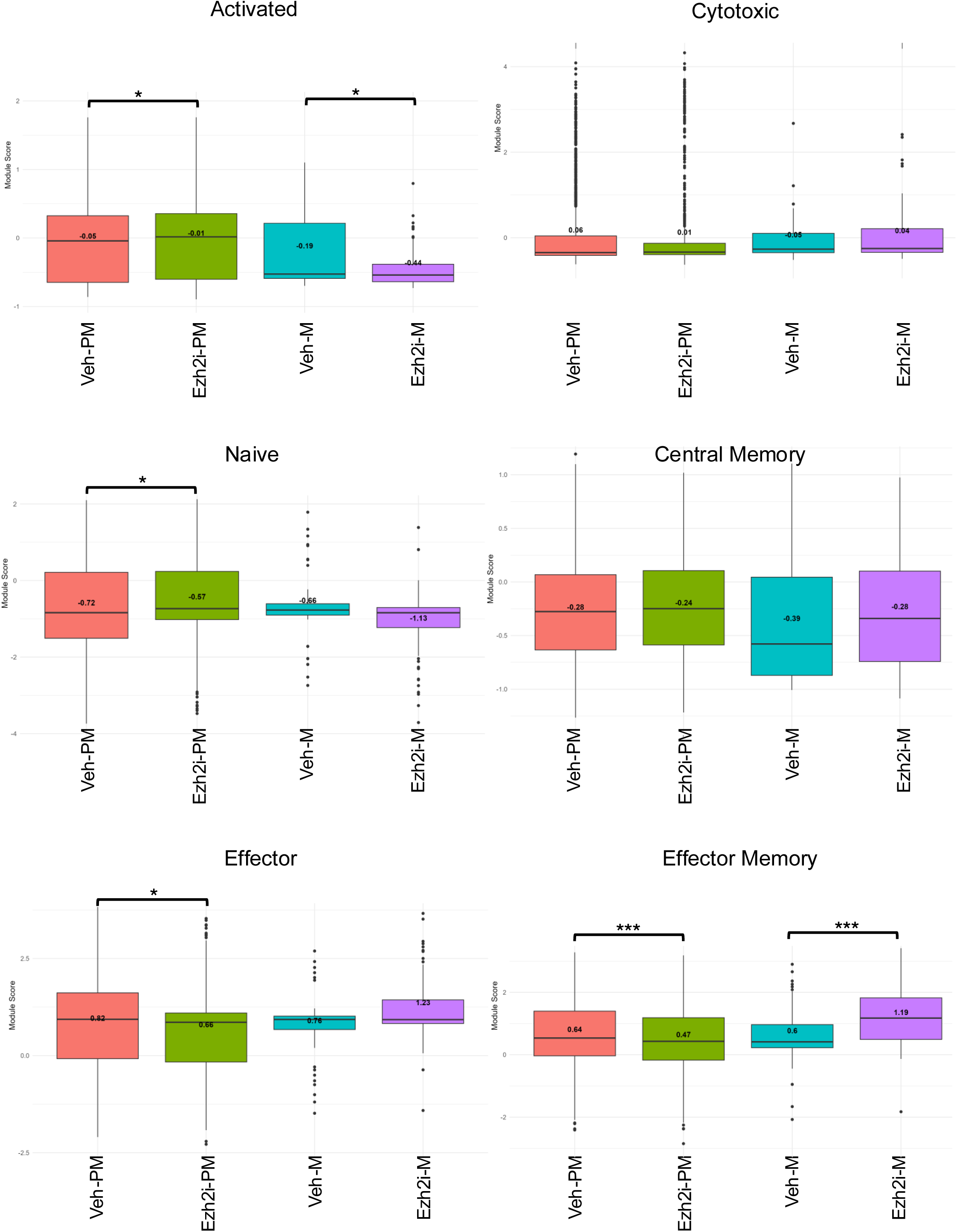
T cell memory states. Box plots of module scores for memory markers defined by the *Mouse Immune Cell Marker Guide* are shown with multiple comparisons across conditions. ns: p > 0.05, *p <= 0.05, **p <= 0.01, ***p <= 0.001, ****p <= 0.0001

**Supplemental Fig. 6.**
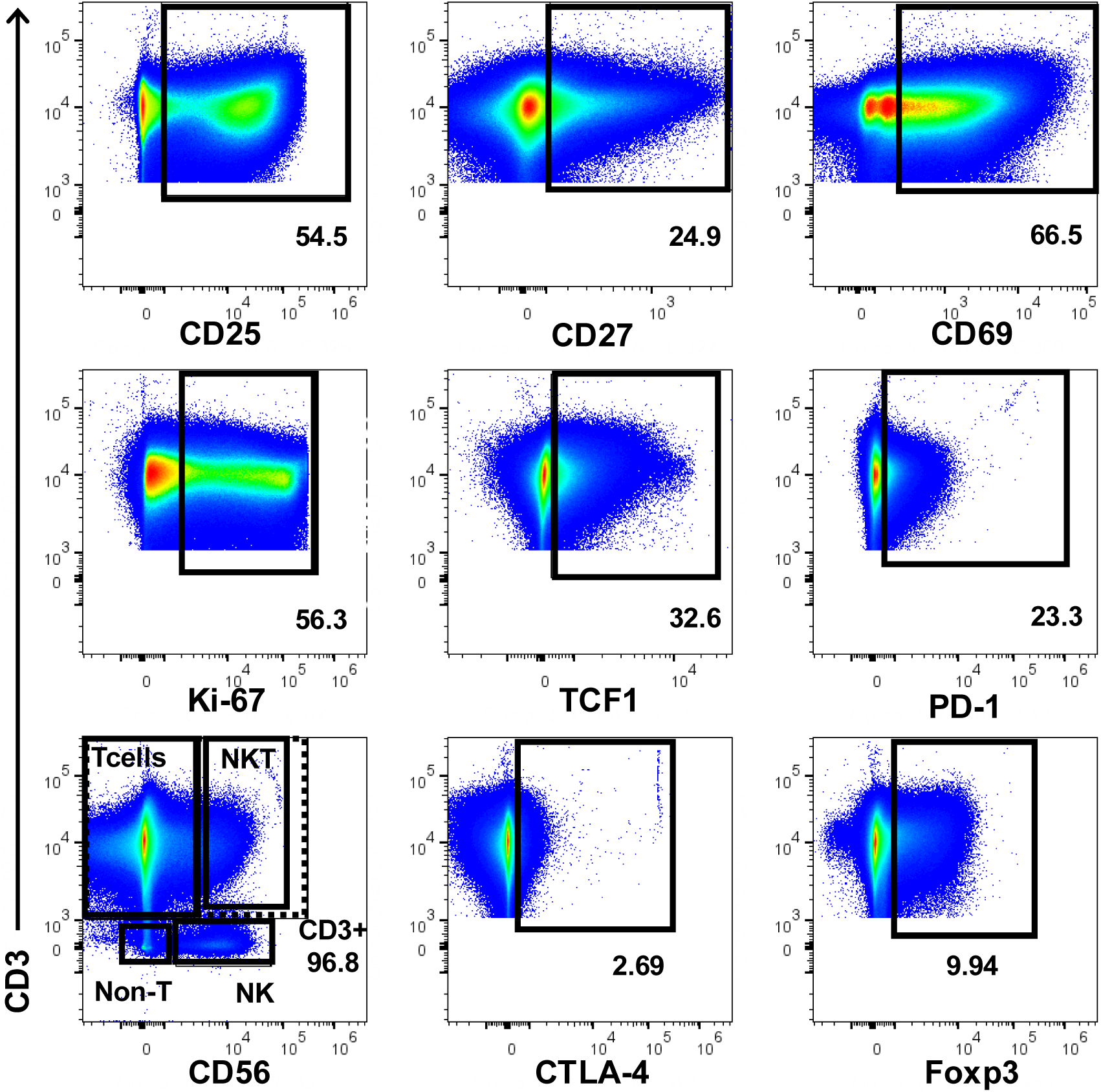
Remaining gating strategies, shown on all-concatentated samples.

## Notes

https://github.com/drplaugher/EZH2-NSCLC-scRNAseq-TILs

### Competing Interest Statement

The authors have declared no competing interest.

### Summary of Updates

manuscript largely updated and revised according to reviewer comments and feedback

